# Spatial biodiversity indicators and a composite index for conservation prioritization in Switzerland

**DOI:** 10.1101/2025.06.10.657334

**Authors:** Antoine Adde, Victor Boussange, Yohann Chauvier-Mendes, Marie-Ange Dahito, Johan Früh, Andrin Gross, Silvia Stofer, Emmanuel Rey, Petra Sieber, Fabian Fopp, Rafael Schouten, Bram Van Moorter, Antoine Guisan, Catherine Graham, Loïc Pellissier, Niklaus E. Zimmermann, Florian Altermatt

## Abstract

Spatially explicit indicators that quantify to which extent landscapes support biodiversity are essential for guiding evidence-based conservation planning. Here, we present a 25-meter resolution dataset for Switzerland, encompassing three key biodiversity indicators— Complementarity, Extinction Risk, and Ecological Connectivity—developed across 17 major taxonomic groups, using habitat suitability maps for approximately 7,500 individual species. These three indicators capture (1) the contribution of landscapes to taxonomic, functional, and phylogenetic diversity, (2) species vulnerability to extinction and (3) ecological connectivity. Outputs are provided for both terrestrial and aquatic realms, including versions adjusted for species richness, and integrated into composite indices. In total, 272 spatial layers were produced and made openly accessible. Relationships among the three indicators were analyzed to assess their distinct contributions, and expert-based validations were performed to evaluate their ecological plausibility and relevance for conservation applications. This dataset provides a robust foundation to support spatial planning and conservation decision-making in Switzerland and can be used as a blueprint for analogue integration in other countries.

## Background & Summary

Spatially explicit biodiversity data support the identification and prioritization of ecologically important areas and are essential for developing effective conservation strategies (Kukkala & Moilanen 2013; Meyer *et al*. 2015; Hochkirch *et al*. 2021). As biodiversity continues to decline globally (IPBES 2019; Giakoumi *et al*. 2024; Keck *et al*. 2025), conservation planning increasingly relies on fine-scale, evidence-based information to guide urgent protection efforts and respond to emerging threats. Such data are also critical for informing land-use planning across sectors such as renewable energy development (Gasparatos *et al*. 2017), transportation infrastructure (Grilo *et al*. 2021), and agriculture (Hoang *et al*. 2023), where competing demands for land often lead to the loss of natural ecosystems and with it the provision of important services (Nick *et al*. 2024). By enabling the assessment of trade-offs and synergies between biodiversity conservation and other land uses, spatial biodiversity data provide a crucial foundation for integrated decision-making (Fastré *et al*. 2021; Hoffmann 2022; Fourchault *et al*. 2024).

Biodiversity’s multifaceted nature calls for multidimensional indices to effectively guide conservation priorities (Devictor *et al*. 2010; Thuiller *et al*. 2014; Chauvier-Mendes *et al*. 2024). No single metric can capture the full complexity of biodiversity, which includes, but is not limited to, species richness, evolutionary distinctiveness, ecological functions and processes (Cadotte *et al*. 2011; Jarzyna & Jetz 2016; Keeley *et al*. 2021). By unifying these diverse yet complementary dimensions within a single analytical prioritization framework, conservation planning can gain a more holistic perspective that accounts not only for species richness but also for species uniqueness and the ecological processes vital in sustaining long-term ecosystem resilience (Mouillot *et al*. 2013; Pollock *et al*. 2017; Beger *et al*. 2022).

International efforts to conserve biodiversity, such as those outlined in the Kunming-Montreal Global Biodiversity Framework, require robust indicators to inform conservation policy at the national scale (Jetz *et al*. 2019; CBD 2022; Eckert *et al*. 2023). As a signatory to the Convention on Biological Diversity, Switzerland formalized its commitment through the Swiss Biodiversity Strategy in 2012 and its corresponding action plan in 2017 (FOEN 2017). The country hosts exceptional biodiversity shaped by its complex geological and climatic history (FOEN 2023). However, this diversity is increasingly threatened by land-use and climate change, habitat fragmentation, pollution, and invasive species, while most national targets for biodiversity conservation remain unmet (FOEN 2023; Klaus 2023). Hence, there is an urgent need for spatially explicit data to guide conservation priorities and support the expansion and integration of protected areas within Switzerland, and all other countries globally. For instance, one key aspect for current and future biodiversity policy is the establishment of a functional ecological infrastructure, which is a network of ecologically important and biodiverse areas (FOEN 2017). Doing so requires adequate information on the spatial distribution and connectivity of key biodiversity variables.

Here, we present a dataset of 272 biodiversity layers at 25-meter resolution to support conservation prioritization in Switzerland. Derived from habitat suitability models for ~7,500 species, along with species trait and occurrence data, the dataset integrates three indicators that together quantify how Swiss landscapes support biodiversity: (1) Complementarity, (2) Extinction Risk, and (3) Ecological Connectivity. The Complementarity Indicator (CI) measures the importance of a pixel based on its contribution to taxonomic, functional, and phylogenetic diversity, as well as its distinct composition of species; the Extinction Risk Indicator (ERI) reflects the importance of a pixel for maintaining species, with emphasis on rare and at-risk species; and the Ecological Connectivity Indicator (ECI) captures the role of a pixel in maintaining connectivity across the landscape. Each indicator was computed independently using tailored methodologies, then rescaled to a common scale to enable their integration into a composite index. Both the individual indicators and the composite index are included in the dataset, supporting flexible approaches to conservation prioritization. Relationships among the three indicators were analyzed to assess their distinct contributions, and expert-based validations were performed to evaluate their ecological plausibility and relevance for conservation applications. The dataset is openly available on Zenodo, and the technical specifications described in this article refer to its first public release (v1.0).

## Methods

### Study area

The study area included all of Switzerland, with a total area of about 41,000 km^2^.

### Input data

#### Species’ habitat suitability maps

Habitat suitability maps for 7,461 individual species (see Appendix S1 for the detailed species list) for the period 1980–2021 were sourced from SDMapCH (v1.3) (Adde et al. in prep), a nationwide raster dataset for Switzerland. This dataset provides modelled habitat suitability maps at a 25-meter resolution, with values ranging from 0 (low suitability) to 100 (high suitability). SDMapCH was developed using the N-SDM modelling pipeline (Adde *et al*. 2023), an end-to-end platform based on a spatially nested hierarchical framework. N-SDM enables multi-level and multi-resolution integration of species and covariate data, effectively addressing challenges such as niche truncation (Guisan *et al*. 2025).

#### Species traits

Functional and dispersal traits of each species were retrieved from TraitCH (Chauvier-Mendes et al. in prep), a comprehensive dataset encompassing ecological traits for over 30,000 species across 17 major taxonomic groups. The dataset includes key ecological trait values, known taxonomic synonyms, and hierarchical taxonomic classifications (family, order, class, and phylum). It also provides information on geographic origin and conservation status according to both the Swiss and global IUCN Red Lists.

#### Species occurrence data

Validated occurrence records for each species, aggregated at a 25-meter resolution and spanning the period 1980–2021, were provided by the Swiss Species Information Center (InfoSpecies) (Dépraz *et al*. 2025).

### Complementarity Indicator (CI)

#### Summary

The CI quantifies the ecological importance of each pixel based on its contribution to taxonomic, functional, and phylogenetic diversity, as well as to the composition of biological communities and ecosystems. It highlights areas that offer unique and complementary biodiversity features, helping to ensure broad ecological representation across species and lineages. CI values range from 0 to 1, with higher values indicating pixels whose loss would disproportionately reduce overall biodiversity representation.

#### Computation

To generate the CI indicator, we used the spatial conservation planning software Zonation 5.0 (Moilanen *et al*. 2022). Based on biodiversity feature inputs, the software ranks pixels of a study region from lowest (0) to highest (1) conservation value. The core principle of Zonation is an iterative removal process: at each step, the pixel contributing least to the overall biodiversity representation is removed, and the values of the remaining pixels are recalculated. This results in a hierarchical prioritization of the landscape, with the most critical pixels for biodiversity conservation ranked highest.

Here, we applied the Core Area Zonation 2 (CAZ2) marginal loss rule algorithm to the set of individual species’ habitat suitability maps of each target taxonomic group (see Taxonomic Output Resolution section). This algorithm gives a particular focus on improving the protection of the least represented species, while maintaining a reasonable level of conservation across all species. As a result, the method ensures strong representation of rare and range-restricted species by safeguarding their core habitats throughout the prioritization process.

Zonation allows users to assign weights to input features to reflect their relative importance in the prioritization. In our case, we incorporated species-level weights based on each species’ phylogenetic and functional uniqueness to give greater importance to those contributing disproportionately to evolutionary history and ecosystem functioning. For each species *i* within taxonomic group *j*, we calculated its functional uniqueness (*FU)* and phylogenetic uniqueness (*PU)* following a previously described method (Grenié *et al*. 2017). Briefly, this method quantifies how functionally or phylogenetically distinct a species is relative to others in the community, based on pairwise trait or phylogenetic distances and species’ relative occurrence. To that end, the functional dendrograms of each taxonomic group were generated based on traits with ≥ 90% species coverage, or > 20% and an imputation error < 0.2 (Soria *et al*. 2021), and the Gower’s distance method (Pavoine *et al*. 2009). Additionally, we constructed each phylogeny with the Open Tree of Life database (Michonneau *et al*. 2016) and assigned branch lengths with the Grafen’s method (Grafen 1989) using the ape R package (Paradis & Schliep 2019). *FU* and *PU* were normalized to a 0–1 range within each taxonomic group to allow comparability and summed to derive a combined uniqueness index: *U*_*i,j*_ = *norm*(*FU*_*i*_)_*j*_ + *norm*(*PU*_*i*_)_*j*_. To avoid assigning a zero weight to any species, cases where *U*_*i,j*_ = 0 were replaced with the minimum non-zero uniqueness value observed within the corresponding taxonomic group.

#### Input data

The CI relied on species-specific habitat suitability maps and trait data. Specifically, we used suitability maps from SDMapCH as the main input features in Zonation, and functional and phylogenetic information from TraitCH and the Open Tree of Life, respectively, to derive species-level uniqueness weights.

### Extinction Risk Indicator (ERI)

#### Summary

At each pixel, the ERI captures the vulnerability of a species group to extinction by estimating the risk of species loss under ecological degradation. It integrates species’ range sizes, occurrences, and conservation statuses to assess the potential impact of local species extirpation on national extinction risk, highlighting areas where conservation efforts would be most effective in preventing irreversible biodiversity loss. In particular, the use of combined range maps with SDM allows for more realistic estimations by accounting for geographical barriers. ERI values range from 0 to 1, with higher values indicating pixels whose degradation would result in disproportionately high extinction risk.

#### Computation

The ERI computation method was adapted from a previously developed metric called the Global Extinction Probability (GEP), originally designed to evaluate how regional species losses impact global species richness (Kuipers *et al*. 2019). The GEP measures the importance of each pixel for conserving a species group by accounting for three key components: the species’ occurrence in the pixel, the extent of its global distribution, and its threat level.

The original GEP formulation considers the global range of each species (i.e., its worldwide distribution). However, global distribution data are not always available, or only at coarse resolution. To address this shortfall, we adapted the approach for regional application, which we refer to as the Regional Extinction Probability (REP). This allowed us to estimate high-resolution extinction probabilities at the Swiss scale, considering each species’ range within the country and, when available, its conservation status in Switzerland. The REP is calculated for a species group *G* at pixel *p* as:

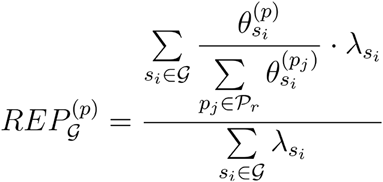

where for each species 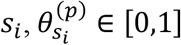 is the occurrence value at pixel *p*, the set 𝒫_*r*_ corresponds to all pixels in the study area (i.e., Switzerland), and 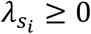 is a weight based on the IUCN conservation status of the species. Weighting schemes for species occurrence 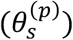 and IUCN conservation status (*λ*_*s*_) are detailed in Text S2.1. In brief, species occurrence values were assigned based on scaled species presence probabilities at each pixel (ranging from 0 for a locally absent species to 1 for a locally present), while conservation status weights followed a linear scale, with higher values given to more threatened species.

To create the Extinction Risk Indicator (ERI), we log-transformed and normalized REP values at the target taxonomic group level (see Taxonomic Output Resolution section). This produced a more balanced distribution and facilitated the comparison and combination of standardized values across indicators. To handle the logarithmic singularity at zero, we replaced all zero values with 10^10−4^, where *n* is one less than the exponent of the smallest non-zero value in the grid. For example, if the smallest non-zero value was at the order of magnitude 10^−3^, then zero values were replaced with 10^−1^. The extinction risk index 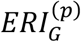 of a group *G* at site *p* is then calculated as:

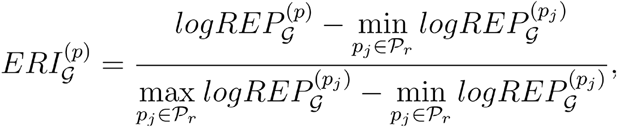

where 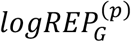 is the logarithmic transformation of 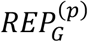, with zero values replaced as described above prior to transformation.

#### Input data

The ERI was based on species-specific habitat suitability maps, occurrence records, and conservation status data. To derive species presence probabilities that account for geographic barriers and historical constraints, habitat suitability maps from SDMapCH were combined with a range mapping algorithm (Fopp et al. in prep). The latter estimated species ranges by clustering validated occurrence records and creating a convex hull and buffer around those clusters of observations. Swiss conservation statuses were obtained from IUCN Red List assessments available in the TraitCH dataset.

### Ecological Connectivity Indicator (ECI)

#### Summary

The ECI evaluates the contribution of each pixel to maintaining ecological connectivity, reflecting its role in enabling species movement between populations. The ECI evaluates the dual function of pixels: as functional habitats hosting dispersing propagules, and as connectors facilitating dispersal between other habitats. ECI values range from 0 to 1, with higher values indicating pixels whose degradation would severely disrupt connectivity, increase fragmentation, and compromise biodiversity persistence and ecosystem function.

#### Computation

The ECI was derived from the Equivalent Connected Habitat (ECH) metric and its partial derivatives (Saura *et al*. 2011; Van Moorter *et al*. 2023a). The ECH is a graph-based metric that can be interpreted as the quality-weighted sum of ecological proximities:

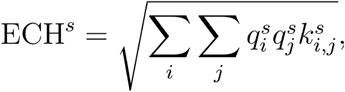

where *ECH*^*s*^ is the ECH value for species *s*, 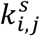 represents the ecological proximity between sites *i* and *j*, and *q*_*i*_^*s*^ is the ecological quality of site *i* for that species. When high-quality sites are either poorly connected or only linked to low-quality sites, the ECH approaches zero (Rubio & Saura 2012; Van Moorter *et al*. 2023b). The ECH is an approximation of the metapopulation capacity, which is numerically more tractable (Song *et al*. 2022). Depending on the species’ movement ecology, ecological proximity between two pixels was computed from their Euclidean distance or from their least-cost path distance derived from a permeability raster, directly obtained from the habitat suitability values. This distance was then scaled by the species-specific average dispersal distance. Ecological quality was directly derived from habitat suitability values. The methods used for computing ecological proximity and distance are detailed in Text S2.2.

The ECI quantifies how a marginal change in habitat quality or permeability at a given pixel affects the ECH for the species. At pixel *i*, these marginal contributions are computed, following Van Moorter et al. (in prep.), as the elasticity of ECH with respect to habitat quality *q*_*i*_ and permeability *p*_*i*_:

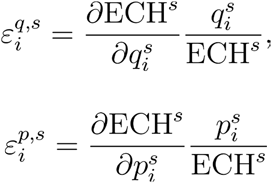

The elasticity with respect to quality 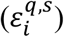 reflects a pixel’s closeness centrality (i.e., how close it is to other high-quality pixels). The elasticity with respect to permeability 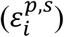 reflects a pixel’s betweenness centrality (i.e., the extent to which it connects high-quality habitats). If a pixel does not contribute to any such connection, its permeability elasticity is zero. These elasticities, calculated with the JAXScape library (https://github.com/vboussange/jaxscape; https://zenodo.org/records/15267704), have been proven reliable proxies for assessing the potential impact of habitat degradation or node removal (Kivimäki *et al*. 2024). Note that 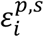 is null when the ecological distance considered for species *s* is the Euclidean distance (see Text S2.2 for details), as the movement of individuals between two functional habitats is assumed to be independent of the intervening pixels. The ECI for pixel *i* and species *s* is defined as the sum of the elasticities, rescaled to range between 0 and 1:

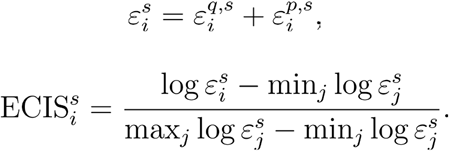

Because the computation of 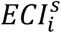 is resource-intensive and our aim was to derive an indicator aggregated at the taxonomic group level (see Taxonomic Output Resolution section), we calculated the ECI for each taxonomic group using a virtual species representative of all species within the group. This virtual species was derived by averaging habitat suitability and dispersal distance across all species within the taxonomic group. In addition, the computational and memory demands for calculating the elasticities increase with the dispersal range. To keep the calculations tractable, we reduced the resolution of the 25-meter suitability maps when computing the partial derivatives 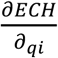 and 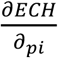, using either 2*δ*^*s*^/10 meters where *δ*^*s*^ is the mean dispersal range for the representative species within the group *s* (see Text S2.2 for details). After computing the derivatives at this coarser resolution, we downscaled the results back to the original 25-meter resolution. Finally, ECI values were log transformed and rescaled between 0 and 1 for harmonization.

#### Input data

The ECI relied on species-specific habitat suitability maps and species traits. Habitat suitability maps from SDMapCH were used to estimate ecological quality and derive permeability rasters. To account for border effects and avoid underestimating connectivity at the boundaries of Switzerland, a buffer zone of 3*δ*^*s*^ meters was included. Values within the buffer were filled using the nearest non-NA value, extending the suitability maps beyond Switzerland’s borders. Species-specific dispersal traits from TraitCH informed the selection of distance types (either least-cost path or Euclidean depending on the species’ movement ecology; see Text S2.2 for details).

### Taxonomic grouping

The three indicators were computed for 17 major taxonomic groups, with a distinction made between terrestrial and aquatic realms within each group (Table 1). “Aquatic” species were defined as those that inhabit or depend on freshwater or wetland environments for key life functions such as foraging or reproduction. This included species occurring in or near water bodies, using aquatic microhabitats, or being strongly associated with riparian or seasonally-wet habitats. This classification was based on habitat usage data, ecological associations, or expert-defined criteria specific to each taxonomic group. All other species were classified as “terrestrial”. It is important to note that the distinction between terrestrial and aquatic species was not always available in the literature, and in such cases, “terrestrial” was applied when the information was unavailable. Species classification resulted in 27 output groups (instead of 34), as some taxonomic groups were exclusively terrestrial or aquatic. In addition, we produced aggregated versions of the indicators for terrestrial (n = 14), aquatic (n = 10), and all groups combined (n = 17), using the maximum pixel value across all relevant groups. This maximum-based aggregation ensures that a pixel is considered important if it is critical for at least one group, offering a conservative and inclusive perspective.

**Table 1.** Summary of the 17 taxonomic groups considered, including the total number of species and their distribution across terrestrial and aquatic realms.

| **Taxonomic Group** | **Number of Species** | **Terrestrial** | **Aquatic** |
| --- | --- | --- | --- |
| Amphibians | 18 | 1 | 17 |
| Bees | 333 | 333 | – |
| Beetles | 605 | 605 | – |
| Birds | 179 | 153 | 26 |
| Bryophytes | 532 | 511 | 21 |
| Butterflies | 717 | 717 | – |
| Dragonflies | 73 | – | 73 |
| Fishes | 37 | – | 37 |
| Fungi | 1449 | 1449 | – |
| Grasshoppers | 92 | 92 | – |
| Lichens | 255 | 255 | – |
| Mammals | 83 | 76 | 7 |
| Mayflies–Stoneflies–Caddisflies | 238 | – | 238 |
| Molluscs | 168 | 122 | 46 |
| Reptiles | 15 | 11 | 4 |
| Spiders | 198 | 198 | – |
| Vascular Plants | 2469 | 2376 | 93 |

### Species richness adjustment

In addition to the uncorrected indicators, we also provide indicators with species richness adjustments. Such adjustments can reduce potential biases arising from areas with high species counts (Safi *et al*. 2011; Zupan *et al*. 2014), particularly in lowland regions. Without these corrections, ecologically valuable areas in Switzerland such as high-altitude or aquatic zones, which tend to host fewer species, may be underrepresented, despite their high conservation importance and potential future relevance under climate change (Adde *et al*. 2024). By accounting for deviations from expected values given local richness, these adjustments provide a more nuanced representation of conservation priorities.

For each indicator, we therefore produced two versions: the raw (unadjusted) values and a version adjusted for species richness. Species richness was calculated as the number of species predicted to be present in each pixel, based on a binarized version of the individual habitat suitability maps. Binarization was performed using species-specific thresholds derived from the maximized True Skill Statistic (maxTSS) criterion (Guisan *et al*. 2017). To correct for the influence of richness, for each output layer, we fitted a Generalized Linear Model (GLM) with quadratic effects, using species richness as predictor (Pardo *et al*. 2017; Chauvier-Mendes *et al*. 2022; Gaüzère *et al*. 2022). The model residuals were retained as richness-adjusted values of our indicators, providing a complementary dimension beyond species richness. Positive residuals indicated higher-than-expected conservation importance, while negative residuals reflected lower-than-expected importance, regardless of species richness. For better integration with the raw versions, all adjusted layers were normalized from 0 to 1.

### Composite index computation

Finally, for the aggregated terrestrial, aquatic, and all-species versions of the indicators, we computed a composite index aimed at identifying areas of high biodiversity value across the three indicators (CI, ERI, and ECI). The composite index was calculated as the pixel-wise mean of the three 0-1 scaled indicator values, providing an integrated metric that reflects multifaceted biodiversity importance. This computation was performed for both the raw (unadjusted) indicator layers and their richness-adjusted versions.

## Data Records

Spatial biodiversity layers for the 17 taxonomic groups, covering the three indicators (CI, ERI, and ECI), their richness-adjusted versions, and composite indices, were compiled into a single dataset (version 1.0), resulting in a total of 272 geospatial layers and a combined file size of approximately 20 GB. These include taxonomic group-specific layers (n = 81), aggregated layers across terrestrial, aquatic, and all species (n = 9), their corresponding richness-adjusted versions (n = 90), and composite indices (n = 6; terrestrial, aquatic, and all-species × two versions). All layers are provided at a 25-meter resolution. The dataset is openly available on Zenodo (Adde et al. 2025).

To illustrate the resulting spatial distribution of the three indicators, we presented example layers (all-species versions), as well as an integrated RGB composite in which each color channel represents one indicator’s relative influence (Figure 1). This visualization highlights areas of convergence and divergence among the indicators, providing an overview of multifaceted priorities for conserving biodiversity across Switzerland. For instance, the Alpine arc with its very specialized high-elevation biota exhibits stronger ERI values, particularly in the central and southern regions. In contrast, the more human-impacted and more fragmented Jura and Plateau regions show stronger CI and ECI contributions. The ECI shows higher values in the valleys, which are natural corridors contributing to connecting habitats. The ERI is generally higher in highlands, where many small-ranged species are found. To explore how these indicators could guide conservation planning, we simulated protected area network scenarios based on their spatial patterns. The corresponding methodology and scenario outputs are detailed in Appendix S3.

**Figure 1.**
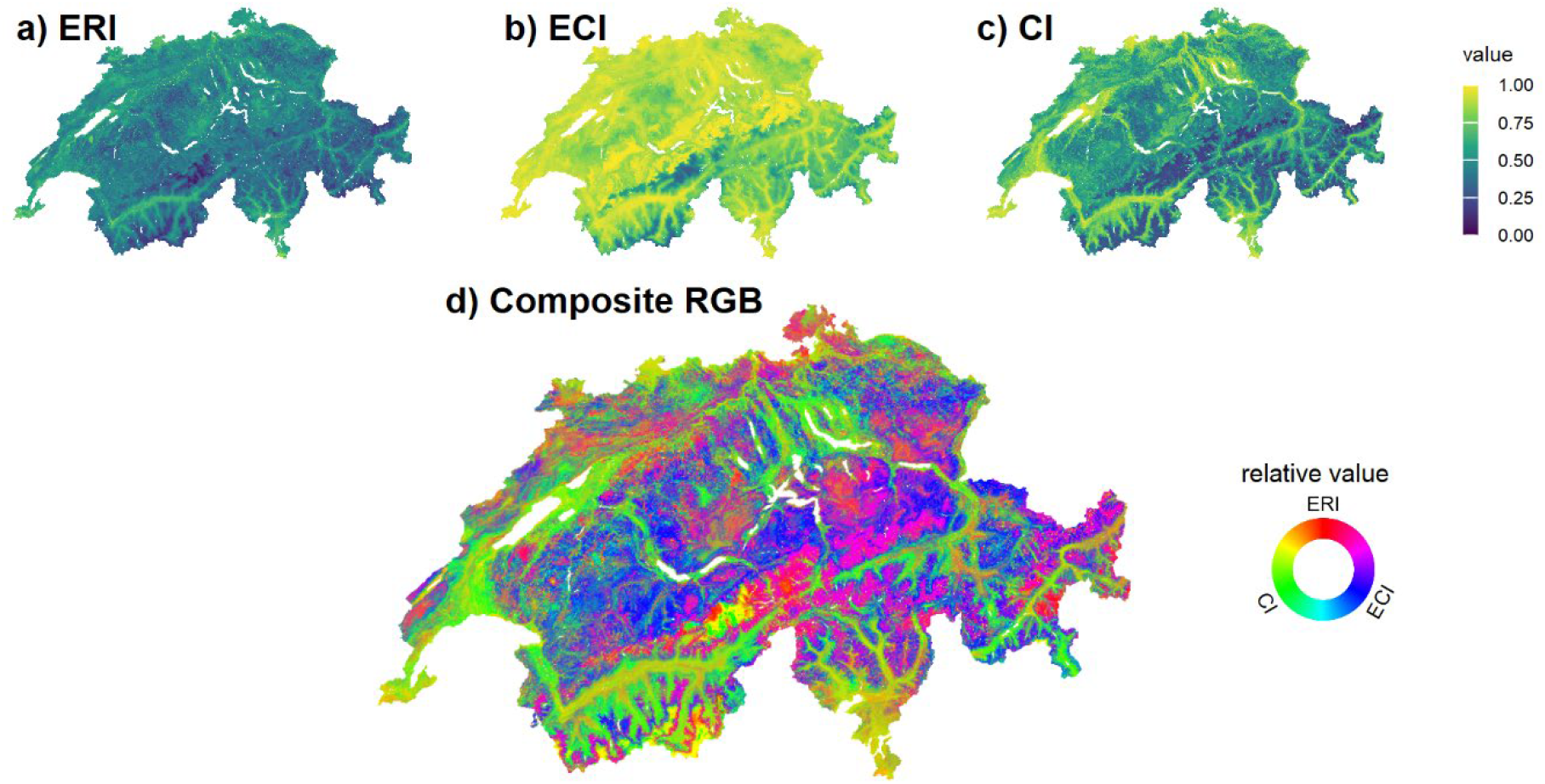
Spatial distribution of the three biodiversity indicators and their composite representation across Switzerland (all-species versions, unadjusted for species richness). (a) Complementarity indicator (CI), (b) extinction risk indicator (ERI), (c) ecological connectivity indicator (ECI), and (d) an RGB composite. In this composite, the green, red, and blue channels represent the normalized relative importance of CI, ERI, and ECI, respectively. For each pixel *(i,j)*, the relative values *x*_*rel*_(*i, j*) are calculated as *x*_*rel*_ (*i, j*) = *x*_*abs*_ (*i, j*) /[*CI*_*abs*_ (*i, j*) + *ERI*_*abs*_ (*i, j*) + *ECI*_*abs*_ (*i, j*)], where *x*_*abs*_ represents the absolute values of either CI, ERI or ECI at pixel *(i,j). CI*_*rel*_, *ERI*_*rel*_ and *ECI*_*rel*_ were stretched to fit the range of 0 to 255 and mapped into an RGB composite. In this composite, pure colors of green represent dominant effects of CI identified across pixels, red for ERI, and blue for ECI. Mixed colors indicate joint effects of two or three drivers.

### Technical Validation

#### Inter-indicator relationships

To evaluate the potential redundancy and distinctiveness of the three indicators (all-species versions), we computed their correlation matrix (Pearson’s r) to quantify the strength of association between them (Figure 2). The results showed a strong relationship between CI and ECI (r = 0.77), indicating that these two indicators captured broadly similar spatial patterns. Moderate R^2^ values were observed between ERI and the other two indicators (r = 0.52 with ECI and r = 0.55 with CI), suggesting that ERI encompasses additional dimensions not fully captured by CI and ECI. These findings indicate that, while some overlap exists, the indicators provide distinct perspectives on how landscapes support biodiversity.

**Figure 2.**
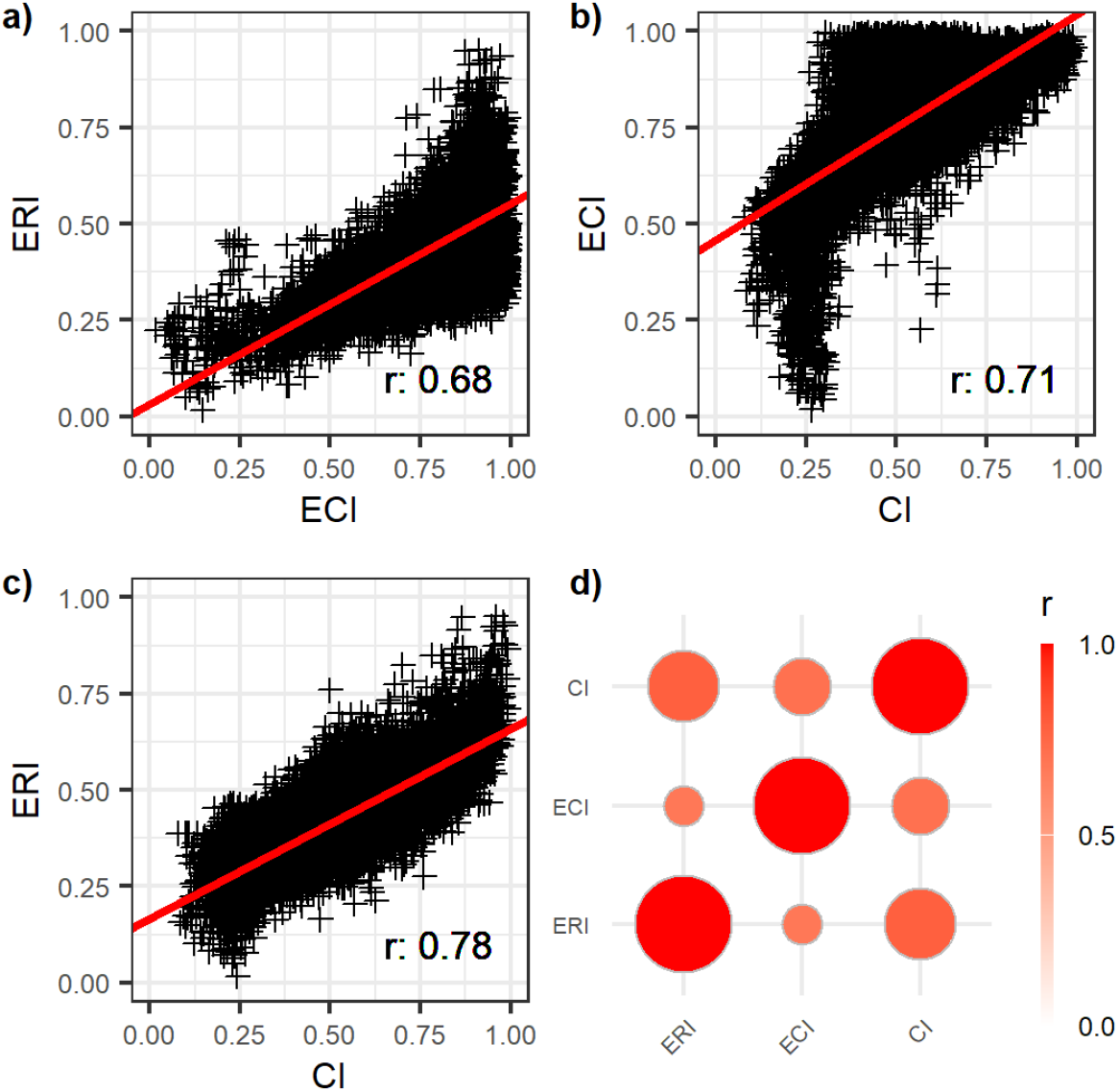
Relationships among the three indicators (all-species versions, unadjusted for species richness, and normalized for comparability) based on simple correlations. (a) Ecological Connectivity Indicator (ECI) as a function of Complementarity Index (CI), (b) Extinction Risk Indicator (ERI) as a function of ECI, and (c) ERI as a function of CI. (d) Correlation matrix of the three indicators. Each plot (a, b, c) shows the fitted regression line (in red) and the coefficient of correlation (r).

### Expert assessment

To evaluate the perceived relevance and utility of the three indicators and their composite index from the perspective of biodiversity experts and conservation professionals, we conducted a qualitative expert-based assessment for the terrestrial, aquatic, and all-species versions of the maps. We submitted the maps to 21 people with expertise in natural history and conservation needs of different taxonomic groups covered, and asked them for evaluation, gathering structured feedback on two main aspects: ecological plausibility and potential value for conservation planning. From the responses received, the indicators were generally considered ecologically sound, with no feedback pointing to implausible patterns. In terms of practical application, while the transition from indicators to implementation guidance remains an open step beyond the scope of this study, the indicators were acknowledged as a relevant first step for data-informed decision-making, particularly in guiding ecological infrastructure and related planning efforts.

### Integrity of output spatial layers

All output layers were validated through an automated procedure that systematically verified compliance with standard specifications, including reference system, spatial resolution, spatial extent, naming convention, and data integrity (number of NA pixels, value range, and data type format). All layers successfully passed the validation process.

## Code Availability

The indicators presented in this manuscript were independently developed by three leading contributors: MAD (ERI), VB (ECI), and YC (CI). The source code for each indicator is available through their respective GitHub repositories:

https://github.com/8Ginette8/CH3Div_complementarity – CI, https://github.com/DahitoMA/CH3Div_extinction – ERI, https://github.com/vboussange/jaxscape – ECI.

## Acknowledgements

This study is part of the SPEED2ZERO Joint Initiative, which received support from the ETH-Board under the Joint Initiatives scheme. The Swiss Species Information Center InfoSpecies (www.infospecies.ch) supplied species occurrence data and expertise on species’ ecology, and we acknowledge their support regarding the dataset. We also thank the experts who evaluated the indicators and provided structured feedback on their ecological plausibility and conservation value.

## Author contributions

**Antoine Adde** co-led the conceptualization of the study and the development and validation of the dataset; led the development of the input habitat suitability maps (SDMapCH); and led the writing of the original draft.

**Victor Boussange** co-led the conceptualization of the study and the development and validation of the dataset; led the development of the ecological connectivity indicator (ECI); and contributed to the writing of the original draft.

**Yohann Chauvier-Mendes** co-led the conceptualization of the study and the development and validation of the dataset; led the development of the complementarity indicator (CI) and the input species trait data (TraitCH); and contributed to the writing of the original draft.

**Marie-Ange Dahito** co-led the conceptualization of the study and the development and validation of the dataset; led the development of the extinction risk indicator (ERI); and contributed to the writing of the original draft.

**Johan Frueh** co-led the conceptualization of the study and the development and validation of the dataset; led the development of the composite index and the protected area network scenarios; and contributed to the writing of the original draft.

**Andrin Gross** contributed to the input species occurrence data and the validation of the dataset.

**Silvia Stofer** led the coordination between the different Swiss data centers for species (InfoSpecies) and contributed to the input species occurrence data and the validation of the dataset.

**Emmanuel Rey** contributed to the input species occurrence data and the validation of the dataset.

**Petra Sieber** contributed to the conceptualization of the composite index; and contributed to the review and editing of the manuscript.

**Fabian Fopp** led the development of the range mapping algorithm; and contributed to the review and editing of the manuscript.

**Rafael Schouten** contributed to the development of the ecological connectivity indicator (ECI), and contributed to the review and editing of the manuscript.

**Bram Van Moorter** contributed to the development of the ecological connectivity indicator (ECI), and contributed to the review and editing of the manuscript.

**Antoine Guisan** co-led the funding acquisition and the development of the input habitat suitability maps (SDMapCH); and contributed to the review and editing of the manuscript.

**Catherine Graham** co-led funding acquisition, supervision, and conceptualization of the study; contributed to the development and validation of the dataset; and contributed to the review and editing of the manuscript.

**Loïc Pellissier** co-led funding acquisition, supervision, and conceptualization of the study; contributed to the development and validation of the dataset; and contributed to the review and editing of the manuscript.

**Niklaus E. Zimmermann** co-led funding acquisition, supervision, and conceptualization of the study; contributed to the development and validation of the dataset; and contributed to the review and editing of the manuscript.

**Florian Altermatt** co-led funding acquisition, supervision, and conceptualization of the study; contributed to the development and validation of the dataset; and contributed to the review and editing of the manuscript.

