## Appendices for "Spatial biodiversity indicators and a composite index for conservation prioritization in Switzerland": _Table of Contents.docx

**– dataset version 1.0 –**

**appendices.zip**

**Table of Contents :**

1. **Detailed species list**
   - File: Appendix S1_species list.csv
   - Description: List of the 7,461 species considered in this study, along with information on the grouping schemes (main taxonomic groups, and aquatic vs terrestrial realms).
2. **Supplementary information on indicator computation**
   - File: Appendix S2_indicator computation.docx
   - Description: Supplementary information on the weighting schemes for species occurrence and IUCN conservation status used for the ERI indicator (Text S2.1) and on the methods used for computing ecological proximity and distance for the ECI indicator (Text S2.2).
3. **Protected areas extension scenarios**
   - File: Appendix S3_protected areas scenarios.docx
   - Description: Simulations of protected areas extension scenarios in Switzerland.
