## Appendices for "Spatial biodiversity indicators and a composite index for conservation prioritization in Switzerland": Appendix S2_indicator computation.docx

### Appendix S2. Supplementary information on indicator computation.

#### Text S2.1. Extinction Risk Indicator (ERI).

##### A) Weighting scheme for species occurrence status.

The weights applied to each species occurrence status (extant to extinct; see Table 1) followed the weighting scheme proposed by (Kuipers *et al.* 2019), which was originally adapted from (Montesino Pouzols *et al.* 2014). To assign occurrence status, we defined categories based on scaled species presence probabilities $\bar{v}$, ranging from 0 to 100. These values were derived from the combination of SDMapCH habitat suitability maps and occurrence records (see section Input data in the main text), processed together using the range mapping algorithm. Scaling enabled for the comparison of values across species when determining occurrence status. The thresholds applied to $\bar{v}$ for assigning occurrence status were designed to be more sensitive at the lower end, where distinguishing true extinction from rare detection is critical. At each pixel, a species *s* is considered to possibly occur if its unscaled presence probability $v$exceeds a species-specific threshold $\tau_{s}$ derived from the maximum True Skill Statistic (maxTSS) criterion (Guisan *et al.* 2017). Since $\tau_{s}$ varies across species, we scale presence probability values such that the species-specific threshold aligns with the central category (i.e., 40), marking the transition to possibly extant status. The scaling transformation is therefore defined as follows:

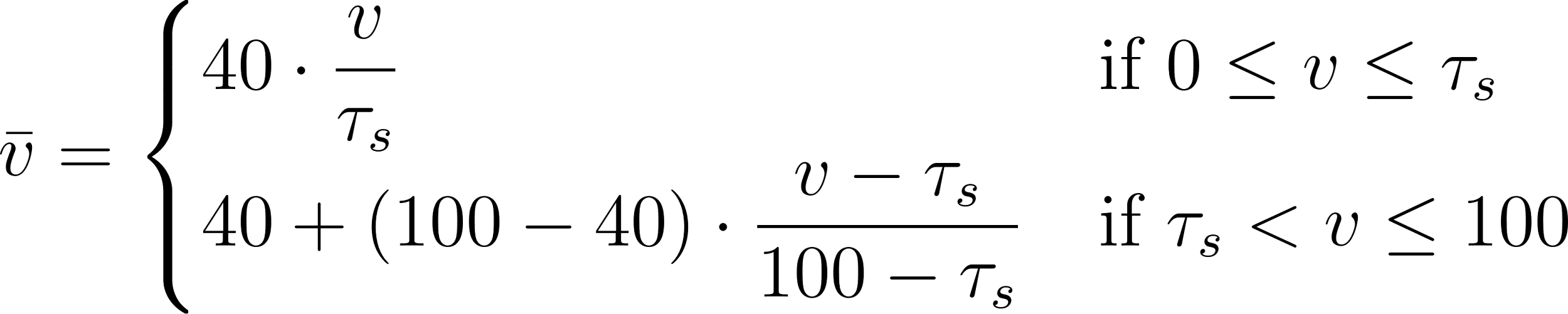
<https://www.codecogs.com/eqnedit.php?latex=%5CBar%7Bv%7D%20%3D%20%5Cbegin%7Bcases%7D%20%2040%20%5Ccdot%20%5Cdfrac%7Bv%7D%7B%5Ctau_s%7D%20%26%20%5Ctext%7Bif%20%7D%200%20%5Cleq%20v%20%5Cleq%20%5Ctau_s%20%5C%5C%20%2040%20%2B%20(100%20-%2040)%20%5Ccdot%20%5Cdfrac%7Bv%20-%20%5Ctau_s%7D%7B100%20-%20%5Ctau_s%7D%20%26%20%5Ctext%7Bif%20%7D%20%5Ctau_s%20%3C%20v%20%5Cleq%20100%20%5Cend%7Bcases%7D#0>

**Table 1.** Weighting scheme for species occurrence $\theta_{s}^{\left( p \right)}$ of species *s* at pixel *p*, based on scaled presence probability values $\bar{v}$*.*

| **Condition on** $\bar{v}$ | **Occurrence Status** | $\theta_{s}^{\left( p \right)}$ |
| --- | --- | --- |
| 80 < $\bar{v}$ ≤ 100 | Present | 1 |
| 60 < $\bar{v}$ ≤ 80 | Probably present | 0.5 |
| 40 < $\bar{v}$ ≤ 60 | Possibly present | 0.5 |
| 20 < $\bar{v}$ ≤ 40 | Possibly absent | 0.1 |
| 10 < $\bar{v}$ ≤ 20 | Probably absent | 0 |
| 0 ≤ $\bar{v}$ ≤ 10 | Absent | 0 |

A) Threat-Level classifications and IUCN conservation status weighting scheme.

In Kuipers et al. (2019), three schemes for quantifying species threat levels were compared (linear, categorical, and logarithmic). Although all three schemes produced similar general patterns, the linear and categorical approaches resulted in closely aligned GEP distributions. In contrast, the logarithmic scheme amplified regional differences, producing more pronounced spatial variation. For consistency with their simulations, we applied the same threat-level classification retained in this previous study, namely the linear scheme detailed in Table 2. We used national conservation status data for Swiss species when available. We made two adjustments to the original threat-level categories: (i) we added the “not evaluated” status, assigning it the same weight as “least concern” and “data deficient,” and (ii) we excluded the obsolete “lower risk” category, which is no longer recognized by the IUCN.

**Table 2.** Weighting scheme associated with the IUCN Red List threat level of species *s*.

| **IUCN Conservation Status** | λₛ |
| --- | --- |
| Critically endangered | 1 |
| Endangered | 0.8 |
| Vulnerable | 0.6 |
| Near threatened | 0.4 |
| Least concern, data deficient, or not evaluated | 0.2 |
| Extinct, extinct in the wild, or regionally extinct | 0 |

#### Text S2.2. Ecological Connectivity Indicator (ECI).

##### A) Ecological proximity.

The calculation of ecological proximity ($k_{i,j}^{s}$*)* was species-specific and was based on a species-specific dispersal kernel ($D^{s}$) and a species-specific ecological distance (see section B below for details on the ecological distance). It was defined as:$k_{i,j}^{s}= D^{s}(d_{i,j}^{s})$, where $d_{i,j}$ was the ecological distance between sites *i* and *j*. We used an inverse exponential function for the dispersal kernel:

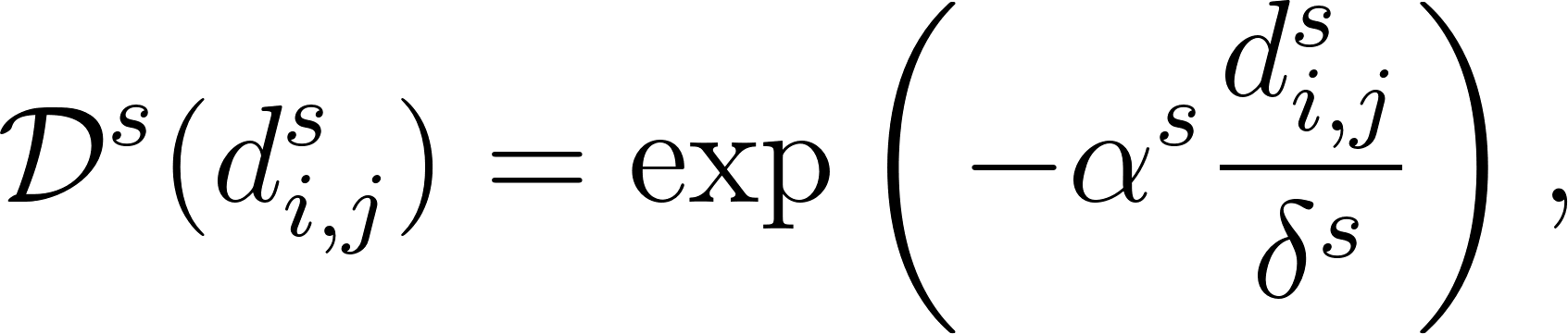
<https://www.codecogs.com/eqnedit.php?latex=%5Cmathcal%7BD%7D%5Es%20(d_%7Bi%2Cj%7D%5Es)%20%3D%20%5Cexp%5Cleft(-%5Calpha%5Es%5Cfrac%7Bd_%7Bi%2Cj%7D%5Es%7D%7B%5Cdelta%5Es%7D%5Cright)%2C#0>

where *δˢ* corresponds to the mean dispersal range for species *s*, expressed in pixel unit, and $\alpha$*ˢ* is a conversion factor to convert $d_{i,j}^{s}$ (expressed in ecological distance) into pixel units. It is calculated as the average ecological distance associated with one pixel:

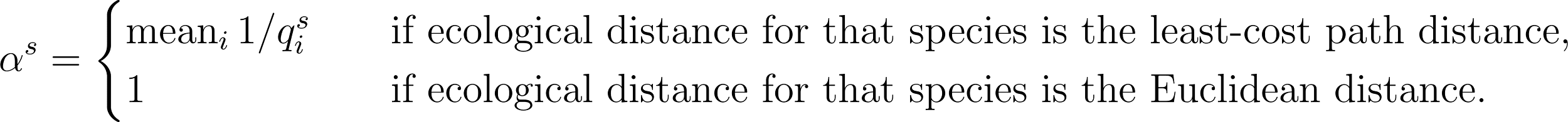
<https://www.codecogs.com/eqnedit.php?latex=%5Calpha%5Es%20%3D%20%5Cbegin%7Bcases%7D%20%5Ctext%7Bmean%7D_i%20%5C%2C%20%7B%201%20%2F%20q_i%5Es%7D%20%26%20%5Cquad%20%5Ctext%7Bif%20ecological%20distance%20for%20that%20species%20is%20the%20least-cost%20path%20distance%7D%2C%5C%5C%20%201%20%26%20%5Cquad%20%5Ctext%7Bif%20ecological%20distance%20for%20that%20species%20is%20the%20Euclidean%20distance%7D.%20%5Cend%7Bcases%7D#0>

The group-specific dispersal distance $\delta^{s}$ were calculated for group S by averaging the respective mean dispersal distance of each species within group S, where the mean dispersal distance for each species was obtained by imputation from the TraitCH dataset (see section Input data in the main text). Specifically, for all taxonomic groups, raw trait information related to species dispersal was identified (Alzate & Onstein 2022) and compiled into a single table, including: average body mass (bm), average body length (bl), maximum body length (mbl), range size (rs), height (h), spore size (ss), spore length (sl), seta maximum length (sml), dispersal ability score (ds), and dispersal distance (dd). Trait availability by taxonomic group is summarized in Table 3. To estimate dispersal distance for all target species, we performed trait imputation using the *missRanger* R package (Mayer & Mayer 2019), incorporating Genus, Family, and Order-level information for each species. In short, the *missRanger* algorithm uses random forest models to iteratively predict missing trait values based on observed data. At each step, imputed values are used to retrain the model, progressively refining the predictions. We parameterized the imputation using the missRanger function and, as recommended in the literature (Stekhoven & Bühlmann 2012; Soria *et al.* 2021), repeated the process 25 times, setting the maximum number of iterations to 20 (maxiter = 20), the number of trees to 100 (num.trees = 100), and enabling predictive mean matching (pmm.k = 3). The latter ensured that imputed values closely matched observed values in the dataset, reducing the likelihood of implausible imputations and preserving the original variance in trait distributions.

**Table 3.** Availability of dispersal-related traits by taxonomic group. Trait abbreviations and units: bm = body mass (g), bl = body length (mm), mbl = maximum body length (mm), rs = range size (km²), h = height (m), ss = spore size (µm), sl = spore length (µm), sml = seta maximum length (mm), ds = dispersal score (unitless), dd = dispersal distance (km).

| **Taxonomic Group** | **bm** | **bl** | **mbl** | **rs** | **h** | **ss** | **sl** | **sml** | **ds** | **dd** |
| --- | --- | --- | --- | --- | --- | --- | --- | --- | --- | --- |
| Amphibians | X | X |  | X |  |  |  |  |  |  |
| Bees | X |  |  | X |  |  |  |  |  |  |
| Beetles |  | X | X | X |  |  |  |  | X |  |
| Birds | X | X |  | X |  |  |  |  |  |  |
| Bryophytes |  |  |  | X |  | X |  | X |  |  |
| Butterflies |  | X |  | X |  |  |  |  | X |  |
| Dragonflies |  | X | X | X |  |  |  |  |  |  |
| Freshwater Fishes |  |  | X | X |  |  |  |  |  |  |
| Fungi |  |  |  | X |  |  | X |  |  |  |
| Grasshoppers |  | X | X | X |  |  |  |  | X |  |
| Lichens |  |  |  | X |  |  |  |  |  |  |
| Mammals | X | X |  | X |  |  |  |  |  | X |
| Mayflies-Stoneflies-Caddisflies |  |  |  | X |  |  |  |  |  |  |
| Molluscs |  | X |  | X |  |  |  |  |  |  |
| Reptiles | X | X |  | X |  |  |  |  |  |  |
| Spiders |  | X |  | X |  |  |  |  | X |  |
| Vascular Plants |  |  |  | X | X |  |  |  |  | X |

##### B) Ecological distance.

The ecological distance $d_{i,j}^{s}$ calculated between sites *i* and *j* captures how a given species perceives the landscape, with its specification depending on the species’ movement ecology (Fletcher Jr *et al.* 2023). As such, we classified species into two broad groups (Table 4):

- For species whose movement between two habitats is significantly influenced by intermediate conditions (i.e., areas or locations that lie between the starting point and destination of a species' movement), we used least-cost path distance. For these species, we used habitat suitability rescaled between 0.05 and 1 as a direct proxy for the site’s permeability to movement.
- For species whose movement was largely unaffected by intermediate conditions, we used Euclidean distance. The calculation of Euclidean distance did not involve the permeability raster.

**Table 4.** Ecological distance type used for each taxonomic group. Least-cost path distance was used for taxa whose movement is constrained by intermediate conditions, while Euclidean distance was used for other taxa which movement is negligibly affected by intermediate sites.

| **Taxon** | **Ecological Distance** |
| --- | --- |
| Mammalls | Least-cost path |
| Reptiles | Least-cost path |
| Freshwater fishes | Least-cost path |
| Molluscs | Least-cost path |
| Bryophytes | Least-cost path |
| Spiders | Least-cost path |
| Insects | Least-cost path |
| Amphibians | Euclidean distance |
| Birds | Euclidean distance |
| Vascular plants | Euclidean distance |
| Fungi | Euclidean distance |
| Lichens | Euclidean distance |
