## Appendices for "Spatial biodiversity indicators and a composite index for conservation prioritization in Switzerland": Appendix S3_protected areas scenarios.docx

**Appendix S3. Protected areas extension scenarios.**

**Text S3.1. Simulating protected areas extension scenarios in Switzerland.**

The aim of this analysis was to simulate the expansion of Switzerland’s protected area network under two scenarios. The first scenario, referred to as the “current” scenario, targets 13.6% protection coverage. This percentage corresponds to the level officially reported by the Federal Office for the Environment (FOEN). However, this level is not directly verifiable using currently available spatial data on protected areas. Therefore, the objective of this scenario was to generate a baseline reference layer that aligns with the reported level. The second scenario, referred to as the “target” scenario, aims to extend protection to 30% of the national territory, reflecting the objective set by international biodiversity conservation agendas.

The simulation used the three biodiversity indicators (CI, ERI, and ECI; all-species versions), as the primary input data. Additional input layers included the existing protected area network (covering 9.4% of Swiss territory, as provided by FOEN, 2024, personal communication), a land impermeability layer representing anthropogenic surfaces (N. Kuelling, 2025, personal communication), and a layer of Swiss lakes retrieved from the [SWECO25](https://zenodo.org/communities/sweco25/records?q=&l=list&p=1&s=10&sort=newest) database.

The method involved several processing steps. First, all lakes larger than 10 km² were excluded (to avoid biasing simulations toward open water areas, which represent a distinct conservation issue), along with impermeable surfaces corresponding to urban or other artificial land cover considered unsuitable for conservation. For each indicator, the remaining pixels were ranked according to their values. A composite index was then calculated as the mean of the three ranked indicators, resulting in a prioritization surface.

The expansion of protected areas was conducted iteratively, beginning from the existing network of protected areas. Pixels for simulated protected areas were added one by one from a pool of potential pixels, starting with those with the highest average rank. At each iteration, potential pixels consisted of pixels that were adjacent to either existing protected areas or previously added pixels. This constraint ensured spatial connectivity and supported the development of realistic expansion zones.

The simulation produced a classification in which each pixel was assigned to one of three categories: (1) unprotected areas, which remained unprotected under the scenario; (2) protected areas, which were already part of the network; and (3) extended protected areas, which were newly added to meet the scenario target. This classification was produced for both the “current” and “target” scenarios, using the composite index in both its species richness-adjusted and unadjusted versions. The output maps, provided at a 100-m resolution, are available in Figure S3.1 and the corresponding R script is provided in Code S3.1.

**Figure S3.1. Simulated protected areas under Current (13%) and Target (30%) scenarios.** GeoTIFF files of the output maps are openly accessible on Zenodo (https://doi.org/10.5281/zenodo.15629783).

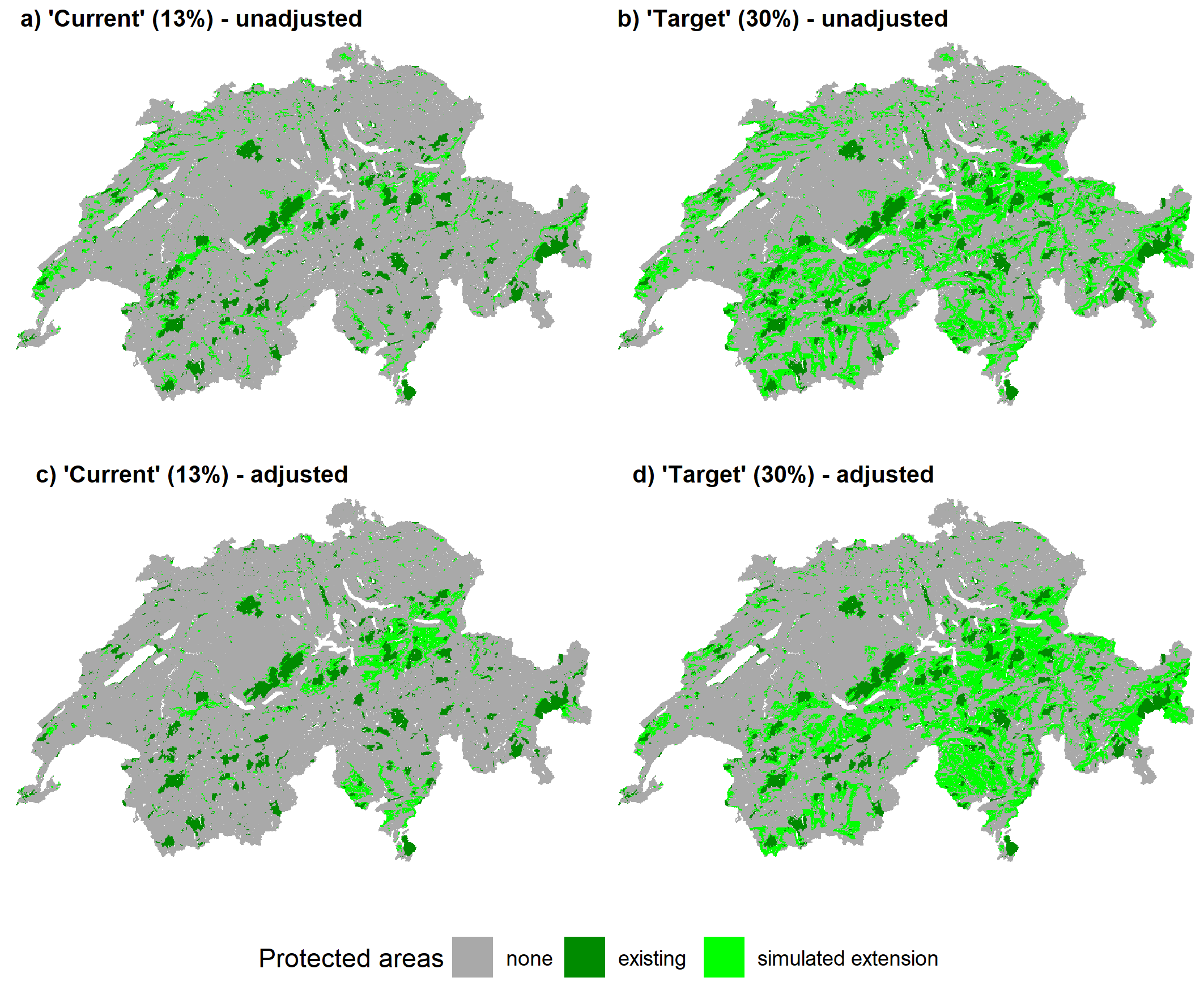

**Code S3.1. R Script for prioritization-based protected Area extension simulation.**

```
library(terra)
library(tidyverse)

# Generate New Protected Areas through pixel selection ####

#load data ####
## mean ranked indices####
r_ECI_CI_ERI_rank_mean <- terra::rast( "CH3Div_Composite_adjusted_All_rankMean.tif")

##existing protected areas (comprehensive: 9.12% of Switzerland)
PAs_comprehensive <- rast( "C:/Users/frueh/Documents/Local Data/Z5_PAs_comprehensive.tif")
#project
PAs_comprehensive <-  terra::project(PAs_comprehensive, "EPSG:2056")

#remove existing PAs from combined layers
r_ECI_CI_ERI_rank_mean_noPAs <- r_ECI_CI_ERI_rank_mean
r_ECI_CI_ERI_rank_mean_noPAs[!is.na(PAs_comprehensive)] <- NA

## impermeability raster ####
r_imperm <- rast("impermeability.tif")
r_imperm_100m <- aggregate(r_imperm, 4)
r_isImperm_100m <- r_imperm_100m > 0

#prepare and align data ####

# resample to 100m (25m too slow and too small for representative conservation areas)
r_ECI_CI_ERI_rank_mean_100m <- aggregate(r_ECI_CI_ERI_rank_mean, 4)
PAs_comprehensive_100m <- aggregate(PAs_comprehensive, 4)

#remove PA surfaces limited to large lake surfaces (align masks)
PAs_comprehensive_100m <- mask(PAs_comprehensive_100m, r_ECI_CI_ERI_rank_mean_100m)

#remove existing PAs surfaces from index mean values
#(these will not be selected, as they are already protected)
r_ECI_CI_ERI_rank_mean_noPAs_100m <- r_ECI_CI_ERI_rank_mean_100m
r_ECI_CI_ERI_rank_mean_noPAs_100m[!is.na(PAs_comprehensive_100m)] <- NA

#count pixels
#determine exact % of comprehensive PAs (this may be different from 9.12% number due to aggregations and lake removals)
pixelNb_existingPAsPerc_100m <- sum(!is.na(values(PAs_comprehensive_100m))) / sum(!is.na(values(r_ECI_CI_ERI_rank_mean_100m)) )
# 0.07353661

pixelNb_existingPAsTotal_100m <- sum(!is.na(values(PAs_comprehensive_100m)))

#pixels needed for reaching exactly 13%
pixelNb_13perc_100m <- floor(pixelNb_existingPAsTotal_100m / pixelNb_existingPAsPerc_100m * (0.13 - pixelNb_existingPAsPerc_100m))

#pixels needed for reaching exactly 30%
pixelNb_30perc_100m <- floor(pixelNb_existingPAsTotal_100m / pixelNb_existingPAsPerc_100m * (0.30 - pixelNb_existingPAsPerc_100m))

#determine starting acceptable cells####
#remove areas outside
# buffer PAs by 1 cell
PAs_comprehensive_100m_buff <- buffer(PAs_comprehensive_100m,100)
#remove existing PAs
PAs_comprehensive_100m_buff[!is.na(PAs_comprehensive_100m)] <- NA
#remove FALSE values (assign NA)
PAs_comprehensive_100m_buff[!PAs_comprehensive_100m_buff] <- NA

#maintain same mask as mean index layer (avoids acceptable pixels generated within large lakes)
PAs_comprehensive_100m_buff <- mask(PAs_comprehensive_100m_buff, r_ECI_CI_ERI_rank_mean_100m)

# remove impermeability
PAs_comprehensive_100m_buff[r_isImperm_100m] <- NA
r_ECI_CI_ERI_rank_mean_noPAs_100m[r_isImperm_100m] <- NA

#get acceptable cell values
acceptableCells <- cells(PAs_comprehensive_100m_buff)

#starting unacceptable cells ####
# (outside boundaries or in PAs)
r_isNA <- is.na(r_ECI_CI_ERI_rank_mean_noPAs_100m)
#remove false values (this is to only get the cell ids of interest)
r_isNA[!r_isNA] <- NA
# get unacceptable cells ids
unacceptableCells <- cells(r_isNA)

#make dataframe with mean ranked index values, and associated cell ids
df_priorConnERisk_100m_mean <- as.data.frame(r_ECI_CI_ERI_rank_mean_noPAs_100m, cells = TRUE)
#order by mean ranked index value
df_priorConnERisk_mean_100m_ordered <- df_priorConnERisk_100m_mean %>% arrange(desc(layer))

#get starting acceptable cells, ordered
acceptableOrdered <- df_priorConnERisk_mean_100m_ordered[df_priorConnERisk_mean_100m_ordered$cell %in% acceptableCells,]

#prepare container for new pixels of protected areas
newPACells <- c()


# FUNCTION: for selecting cells ####
##C_RATE: pixel rate (how many pixels are placed simultaneously per iteration)
##C_min : starting value (allows to extend results from the 13% run to achieve 30%, rather than starting over)
##C_max: value (to reach 13% or 30% of pixels)

func_selectNewPAs <- function(C_RATE, C_min, C_max, newPACells = newPACells,
                              acceptableCells = acceptableCells, acceptableOrdered = acceptableOrdered, unacceptableCells = unacceptableCells){

  #iterate to get all new cells required to fill 13% or 30%
  for(i in seq(from = C_min, to = C_max, by = C_RATE)){

    #show progress
    print(Sys.time())
    print(paste0(i, "-", i+C_RATE," cells"))
    #subset acceptable from all ordered cells
    acceptableOrdered <- df_priorConnERisk_mean_100m_ordered[df_priorConnERisk_mean_100m_ordered$cell %in% acceptableCells,]
    #take the highest valued C_RATE cells
    #(or the remaining number, if not divisible by C_RATE)
    if(C_max-i >= C_RATE){

      #get highest values C_RATE cells
      newCells <- acceptableOrdered$cell[1:C_RATE]

    }else{

      #get remaining number of cells
      newCells <- acceptableOrdered$cell[1:(C_max - i + 1)] #+1 otherwise last pixel is missing

    }

    #add new chosen cells
    newPACells <- c(newPACells, newCells)

    #remove newly chosen cells from acceptableCells
    acceptableCells <- acceptableCells[!acceptableCells %in% newCells]

    #remove new cells from acceptable
    acceptableOrdered <- acceptableOrdered[(C_RATE+1):nrow(acceptableOrdered),]

    #include new adjacent cells in acceptable
    adjacentCells <- unique(as.numeric(terra::adjacent(r_ECI_CI_ERI_rank_mean_noPAs_100m$layer, newCells)[,2:4]))
    #only keep those that are not unacceptable AND not in new cells
    acceptableAdjacent <- adjacentCells[!(adjacentCells %in% unacceptableCells) & !(adjacentCells %in% newCells)]
    acceptableCells <- c(acceptableCells, acceptableAdjacent )

    #make new cells unacceptable (to not sample them again)
    unacceptableCells <- c(unacceptableCells, newCells)
    #

  }
  return(list(newPACells = newPACells, acceptableCells = acceptableCells, acceptableOrdered = acceptableOrdered, unacceptableCells = unacceptableCells))
}

#determine parameters (starting number of pixels, target number of pixels, rate of pixels)
#C_RATE: pixel rate (how many pixels are placed simultaneously per iteration)
#C_min : starting value (allows to extend results from the 13% run to achieve 30%, rather than starting over)
#C_max: value (to reach 13% or 30% of pixels)

#launch first round to reach 13%
results_13perc <- func_selectNewPAs(C_RATE = 100, C_min = 0, C_max = pixelNb_13perc_100m,
                                    newPACells = c(),
                                    acceptableCells = acceptableCells,
                                    acceptableOrdered = acceptableOrdered,
                                    unacceptableCells = unacceptableCells)

#launch second round, using results from 13%, to reach 30%
results_30perc <- func_selectNewPAs(C_RATE = 100, C_min = pixelNb_13perc_100m, C_max = pixelNb_30perc_100m,
                                    newPACells = results_13perc$newPACells,
                                    acceptableCells =  results_13perc$acceptableCells,
                                    acceptableOrdered = results_13perc$acceptableOrdered,
                                    unacceptableCells = results_13perc$unacceptableCells)

# generate rasters for new PAs
#prepare empty raster with proper masks
r_newPAs_13perc <- rast(resolution = 100, crs = "EPSG:2056", extent = ext( r_ECI_CI_ERI_rank_mean_noPAs_100m$layer) )
r_newPAs_13perc[1:ncell(r_newPAs_13perc)] <- 0
r_newPAs_13perc <- mask(r_newPAs_13perc,r_ECI_CI_ERI_rank_mean_100m$layer )
#give 1 to the determined new PA cells
r_newPAs_13perc[results_13perc$newPACells] <- 1
#give 2 to existing PAs
r_newPAs_13perc[PAs_comprehensive_100m == 1] <- 2

# terra::writeRaster(r_newPAs_13perc, "newPAs_13perc.tif")

#plot areas for new PAs
#prepare empty raster with proper masks
r_newPAs_30perc <- rast(resolution = 100, crs = "EPSG:2056", extent = ext( r_ECI_CI_ERI_rank_mean_noPAs_100m$layer) )
r_newPAs_30perc[1:ncell(r_newPAs_30perc)] <- 0
r_newPAs_30perc <- mask(r_newPAs_30perc,r_ECI_CI_ERI_rank_mean_100m$layer )
#give 1 to the determined new PA cells
r_newPAs_30perc[results_30perc$newPACells] <- 1
#give 2 to existing PAs
r_newPAs_30perc[PAs_comprehensive_100m == 1] <- 2

# terra::writeRaster(r_newPAs_30perc, "newPAs_30perc.tif")
```
